# Conjugation mediates large-scale chromosomal transfer in *Streptomyces* driving diversification of antibiotic biosynthetic gene clusters

**DOI:** 10.1101/2024.05.27.596038

**Authors:** Caroline Choufa, Pauline Gascht, Hugo Leblond, Anthony Gauthier, Michiel Vos, Cyril Bontemps, Pierre Leblond

## Abstract

*Streptomyces* are ubiquitous soil dwelling bacteria of special importance as a source of metabolites used in human and veterinary medicine, agronomy and industry. Conjugation is the main mechanism of *Streptomyces* Horizontal Gene Transfer, and this process has long been known to be accompanied by mobilization of chromosomal DNA. However, the magnitude of DNA transfer, or the localization of acquired DNA across their linear chromosome, has remained undetermined. We here show that conjugative crossings in sympatric strains of *Streptomyces* result in the large-scale, genome-wide distributed replacement of up to one third of the recipient chromosome, a phenomenon for which we propose the name ‘*Streptomyces* Chromosomal Transfer’ (SCT). Such chromosome blending results in the acquisition, loss and hybridization of Specialized Metabolite Biosynthetic Gene Clusters, leading to a novel metabolic arsenal in exconjugant offspring. Harnessing conjugation-mediated SMBGC diversification holds great promise in the discovery of new bioactive compounds including antibiotics.

## Introduction

Horizontal gene transfer (HGT) is a major driver of the diversification and adaptation of bacteria^1^. Conjugation, where gene transfer is dependent on cell-cell contact, is one of the main mechanisms of uptake of foreign DNA^2–4^. Conjugation is mostly mediated by autonomous Mobile Genetic Elements (MGEs) that ensure their own transfer from the donor to the recipient strain, either remaining extrachromosomal (plasmids) or integrating into the recipient’s chromosome (Integrative and Conjugative Elements, ICEs). Early work using the F-factor of *Escherichia coli* demonstrated that conjugation can also result in the transfer of chromosomal DNA flanking the integration site (Hfr^5^). A variety of other conjugation systems leading to large-scale chromosomal DNA transfer has since been described, including in *Mycobacterium smegmatis*^6^ and in *Mycoplasma agalactiae*^7^.

Pioneering work by Sir David A. Hopwood revealed that recombination in *Streptomyces coelicolor* A3(2) was stimulated by so-called ‘Fertility Factors’, a feature which was exploited to produce the first genetic maps in *Streptomyces*^8^. These fertility factors were further characterized as conjugative plasmids or AICEs (Actinomycete ICEs; the most prevalent MGEs in *Streptomyces* genomes^9–11)^. AICEs and conjugative plasmids in *Streptomyces* utilize a fundamentally different system for self-transfer compared to canonical conjugative elements. During transfer, AICEs excise from the donor chromosome, replicate and use the DNA translocase TraB, forming a conjugation pore, to self-transfer as dsDNA between hyphal compartments after which they reintegrate in both donor and recipient target insertion sites^12,13^.

Transfer of large chromosomal regions accompanying that of conjugative elements has long been suspected in *Streptomyces* based on the finding that recombinants often inherited distant markers from a parental donor strain^8^. We here quantify the extent of conjugal transfer of chromosomal DNA in *Streptomyces* via mating assays coupled to the resequencing of transconjugant genomes. To do so, we employed pairs of strains with an overall genomic nucleotide similarity of 98.7% and 99% respectively within each pair; high enough to allow efficient homologous recombination, but divergent enough to enable *in silico* identification of donor DNA in recipient genomes. This approach allowed us, for the first time, to elucidate tract length and distribution of transferred chromosomal- and MGE DNAs in *Streptomyces*. Our experiments also provide an in-depth window into the acquisition and recombination of specialized metabolite biosynthesis gene clusters (SMBGCs)^14,15^ demonstrating that HGT greatly accelerates both the acquisition of novel SMBGCs^16–18^ and their rapid evolution by their recombination.

## Main

### Streptomyces Chromosome Transfer (SCT): large-scale conjugative transfer of chromosomal regions in Streptomyces

To decipher how- and to what extent conjugative elements promotes the transfer of the chromosomal DNA in *Streptomyces*, we performed mating experiments with closely related strains isolated from the same sympatric soil population^19^. These strains that harbored a linear chromosome of ca. 12 Mb were previously shown to contain a great diversity of MGEs, including AICEs capable of effective transfer between strains^11^. Two strain couples (couple A: S1A1-3* x S1D4-23*; couple B: RLB1-9* x SID4-23*; Figure 1 and S1, Table S1) able to reciprocally conjugate were selected for use in this study. Among these three strains, only RLB1-9* possessed plasmids (Table S1). Each strain was chromosomally labeled with different antibiotic resistance genes (labelled strain*^/^**, Table S1) and recombinants selected after mating on plates containing both antibiotics. Mating assays produced recombinants at a frequency of ∼10^-5^-10^-6^, which is in the same range as previously reported for different *Streptomyces* species and conjugative factors^20–23^. Twelve recombinants were picked for each couple and their genomes were sequenced at high coverage (100x). The sequence for each recombinant strain was deduced from the alignment of sequencing reads on the parental chromosomes and SNP analysis was used to identify the parental origin of each recombinant region.

**Figure 1:**
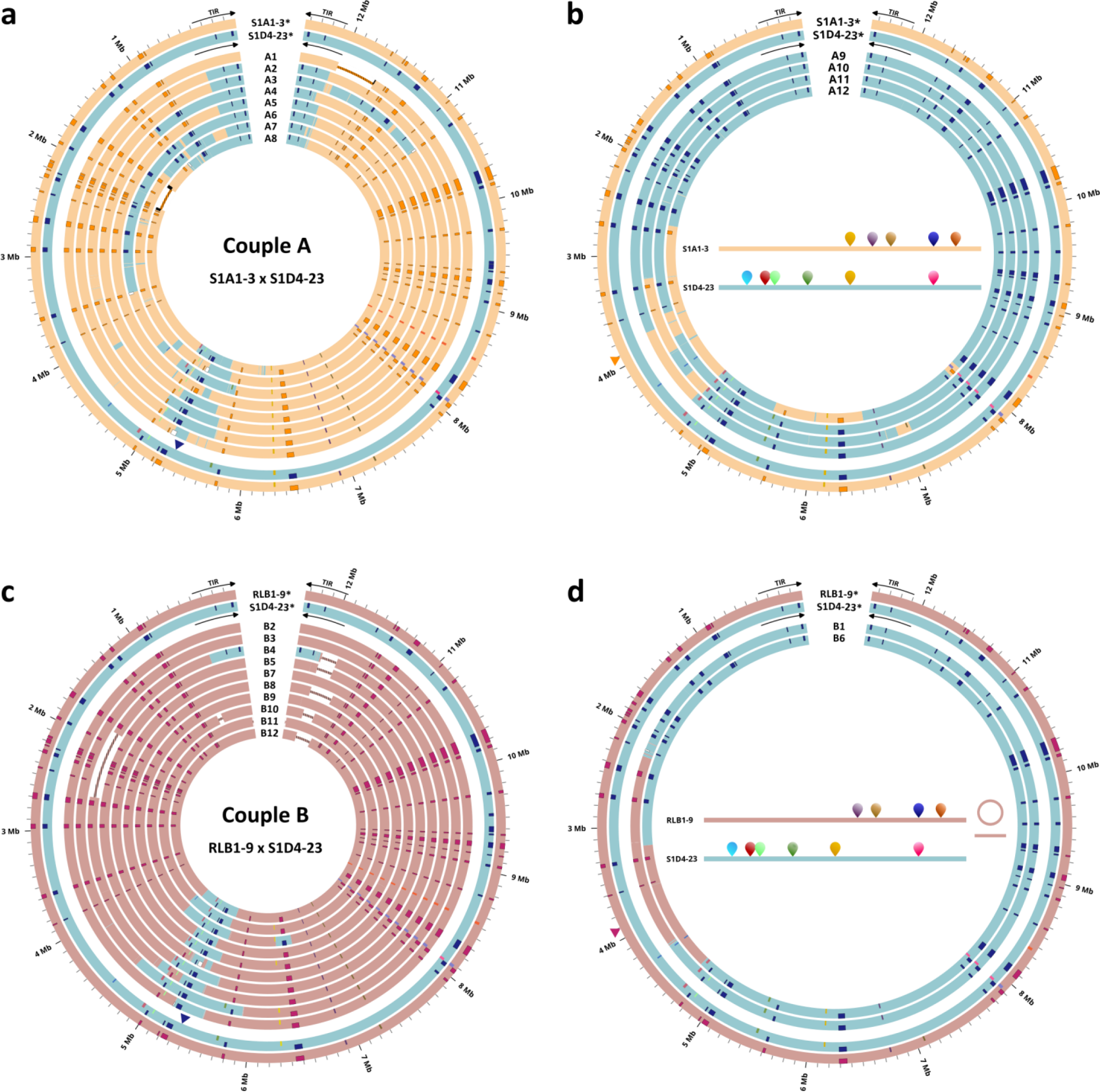
Chromosomal DNA acquisition in recombinant strains. The linear chromosomes of the parental and recombinant strains for mating couple A (S1A1-3* x S2D4-23*) and B (RLB1-9* x S1D4-23*) are represented as open concentric circles (Circos). **a**, recombinants A1 to A8 with a S1A1-3* genomic background. **b**, recombinants A9 to A12 with a S1D4- 23* background. **c**, B2-B5 and B7-B12 recombinants with a RLB1-9* genomic background. **d**, recombinants B1 and B6 with a S1D4-23* genomic background. For each couple, the parental chromosomes correspond to the two outer circles (S1A1-3*, S1D4-23*, RLB1-9*), and the recombinants to the inner circles. Terminal Inverted Repeats (TIRs) symbolized by arrows are shown on the external ring for the parents only. AICEs are represented by rectangles on the lower edge of the concentric circles. SMBGCs are represented by rectangles on the upper edge of the concentric circles. Chimeric and hybrid (rearranged) SMBGCs are colored white and black, respectively. Large deletions are represented by a hatched line. All recombinants have inherited the donor chromosomal selection marker (colored triangle). Each locus (including the selected antibiotic resistance gene) was positioned relative to the chromosomal positions of the recipient genome. In panels **b** and **d**, AICE/plasmid content is represented inside the Circos scheme.

For couple A, eight recombinants had a S1A1-3* genomic background (Figure 1a) and four a S1D4- 23* background (Figure 1b). For couple B, ten recombinants had a RLB1-9* genomic background (Figure 1c) and two a S1D4-23* background (Figure 1d). None of the recombinants was identical to each other. The sizes of donor fragments in recombinant genomes varied from hundreds of kilobases down to extremely short stretches consisting of just a few nucleotides (Figure 2 and S2). As expected following double antibiotic selection, all recipient strains acquired a DNA fragment harboring the antibiotic resistance of the donor strain. For couple A, recombinants acquired 3-52 fragments of the donor DNA with an average total size of 1,825 kb (192 kb - 4,713 kb) representing 1.5% to 37.8% of total parental genome size (Figure 2, Table S2). For couple B, 1 to 13 fragments were acquired with an average total size of 994 kb (255 kb – 2,197 kb) representing 2% to 17.5 % of the total parental genome size. Overall, each recombinant acquired at least one large fragment (size 152 kb -2,020 kb), frequently corresponding to that containing the selected resistance from the donor. In one third of the recombinants (5 out of 24) the second largest chromosomal fragment acquired (after the fragment containing the selection marker) corresponded to the end of the chromosome, including the long terminal inverted repeat (TIR, sizes from 303 kb to 408 kb^24^) capped by the telomere (Table S2). This phenomenon is particularly striking in couple A, where seven S1A1-3* background recombinants acquired both ends of the S1D4-23* chromosome (Figure 1). None of the four recombinants analyzed in the other direction acquired the chromosome ends of the donor (S1A1-3*). For couple B only one TIR replacement was observed for the recombinant B4 (Figure 1c). Smaller fragments, defined as shorter than the median value of the recombined fragments (*i.e.*, 4.9 kb), were predominantly located close to large acquired regions, resulting in “microheterogeneity” hotspots (Figure 2 and S2). In addition to DNA acquisitions, deletions up to several hundreds of kilobases (Figure 1) were observed in almost a third of the recombinants but these occurred independently of DNA acquisitions.

**Figure 2:**
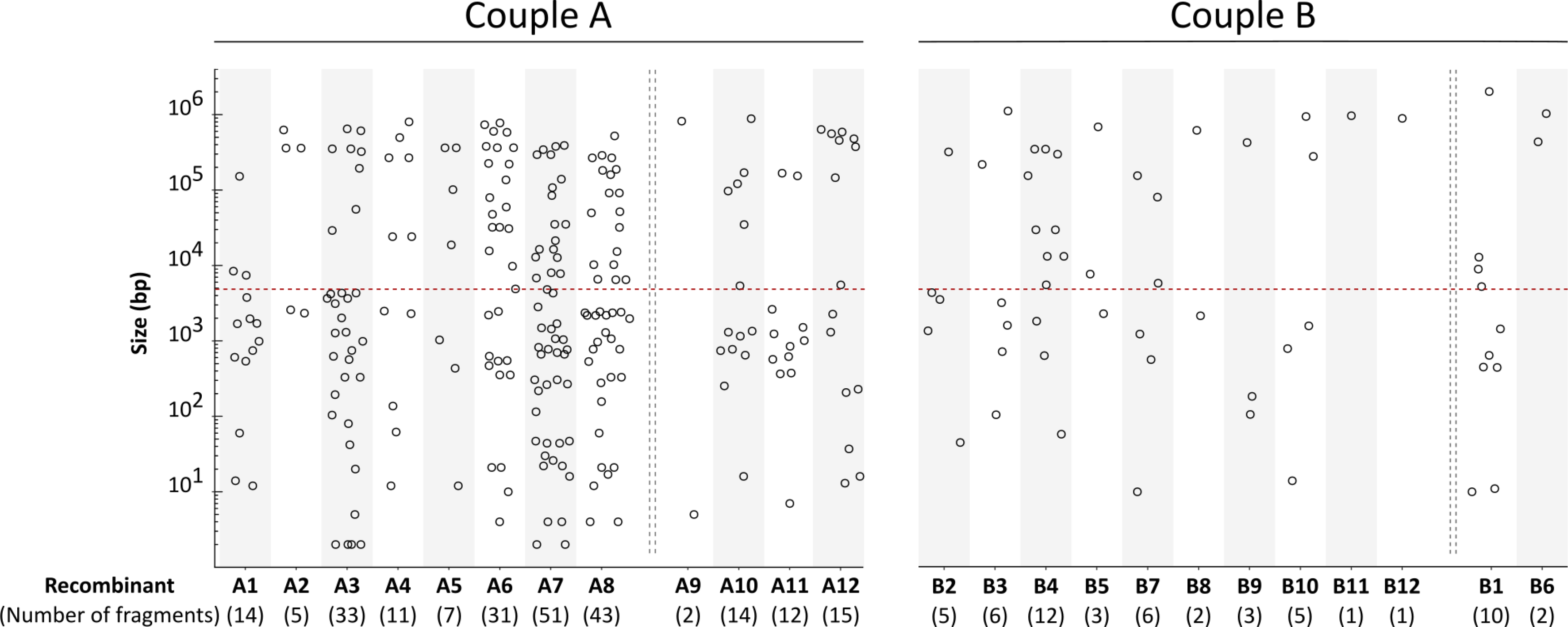
Number and size of incorporated DNA fragments in recombinant progeny. Size length distribution of incorporated donor DNA in all recombinants obtained for couple A and B (A1 to B12). The number of acquired fragments for each recombinant is indicated in brackets. The different recipient genomic backgrounds are separated by a vertical double dotted line. The dotted red horizontal line at 4.9 kb represents the median length of the recombinant DNA fragments.

#### AICEs drive chromosomal mobilization

All 24 recombinants but one acquired one or several AICEs. Eleven AICEs, all possessing the three key functions (*i.e.* integration, replication, transfer) enabling their autonomous mobilization, were present in the parental strains. Eight out of 11 were transferred in the recombinants. Several types of AICE acquisition were observed. Self-transfers of 3 AICEs (AICE05, 11, 13, Figure 3b), i.e. the result of AICE-mediated excision and integration (evidenced by AICE flanking sequences from the recipient strain), was observed in 19 out of 24 recombinants (Figure 3b). In one case, the self-transfer of two AICEs was observed (recombinant B2, Figure 3b). In 14 recombinants, AICEs were acquired not by self-transfer but likely through homologous recombination events with the chromosomal regions flanking the elements. In nine of these recombinants this non-autonomous acquisition occurred along with self-transferred AICEs; in four cases (A2, A8, B9, B12), there is no evidence of AICE self-transfer and/or integration. In recombinant A2, the acquired AICE (AICE05) was flanked by chromosomal DNA of the donor on only one side revealing the mobilization of DNA on only one side of the element (mono-directional mobilization^8^). Interestingly, two recombinants (A3 and B11) were found to have acquired a chromosomal fragment which harbored an AICE in the donor but not in the recipient, possibly because the AICE was subsequently lost during the mobilization process by the defect of the integration step in the recipient chromosome.

**Figure 3:**
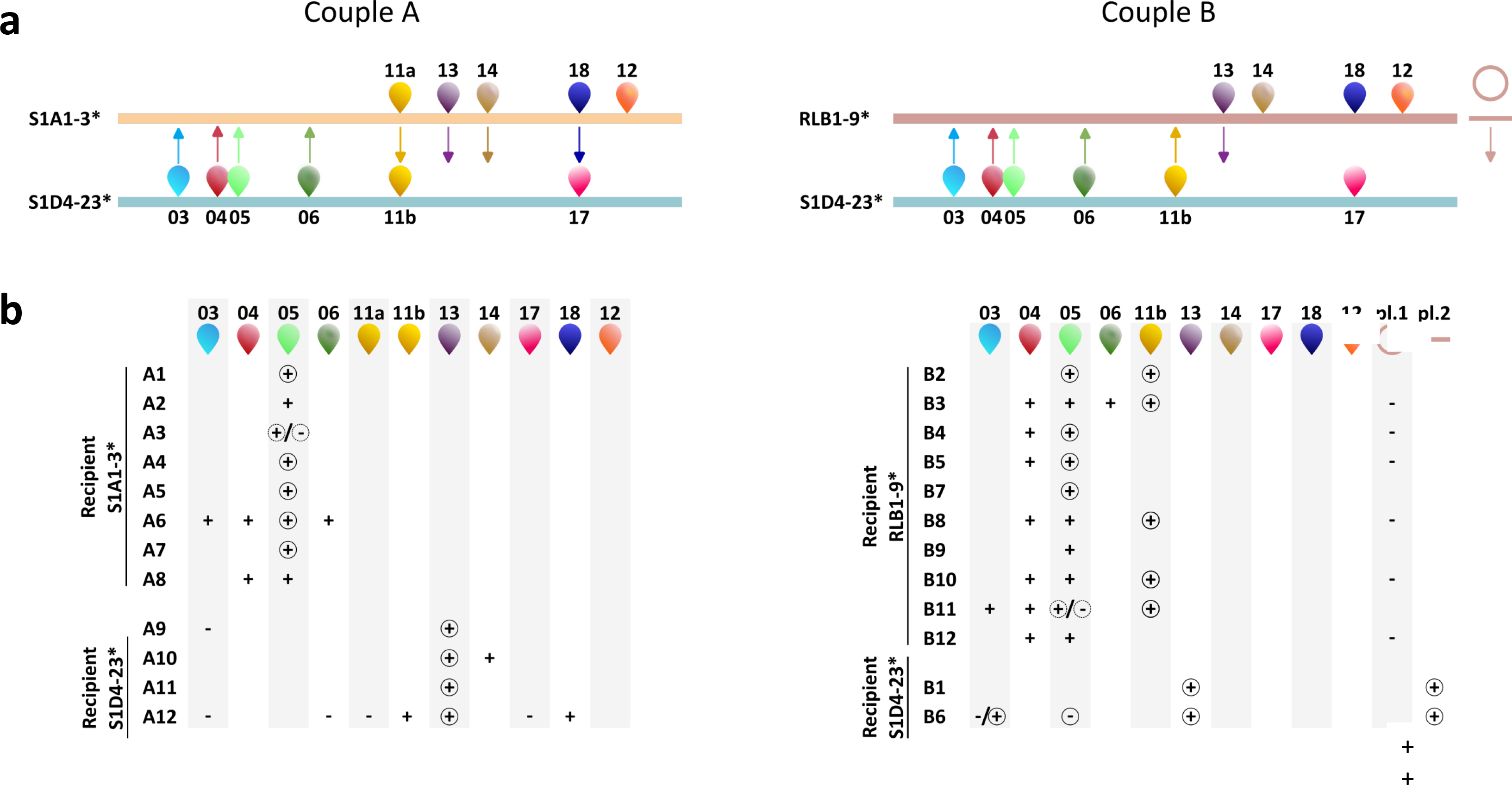
MGE transfer in recombinants. **a**, shows AICE content of the parental strains. AICE mobility is indicated by colored arrows. The mobility of AICE11a and AICE18 (couple A) results in the replacement of AICE11b and AICE17, respectively, at the same insertion site; AICE11a and 11b are different elements but belong to the same family (same color code). **b**, describes the acquired (+)/lost (-) AICEs for each recombinant of the two couples A and B. When the AICE is self-transferred, the + symbol is circled. In recombinants A3 and B11, the +/- circled by a dotted line symbolizes the probable loss (by self-excision) of AICE05 during transfer either in the donor or the recipient. Note that the loss of AICE05 in recombinant B6 probably results from its excision not followed by reintegration in its original chromosome. AICE03 in B6 was probably lost by replacement of the homologous region of the donor devoid of AICE03, but reintegrated at its target site after DNA acquisition. The plasmids pRLB1-9.1 (circular) and pRLB1-9.2 (linear) are represented by a circle and a line, respectively.

During the conjugation process, gain or loss of plasmids in the recombinants was also observed. In couple B (the only couple involving plasmids) the acquisition concerned the linear plasmid pRLB1-9.2 in recombinants B1 and B6. Loss of the circular plasmid pRLB1-9.1 was observed in two thirds of the RLB1-9* recombinants (Figure 3).

### Large-scale recombination of Specialized Metabolite Biosynthetic Gene Clusters (SMBGCs)

Specialized metabolite biosynthesis gene clusters (SMBGCs) are large genomic regions that encompass both biosynthesis genes together with regulatory and resistance genes (in case of toxic activity such as antibiotics). Conjugative HGT was found to result in profound reassortment of parental SMBGCs in recombinants. Twenty-two out of 24 recombinants differed in SMBGC content relative to the two parental strains which contain between 36 to 40 SMBGCs each (Table 1, Figure S3). As most SMBGCs were common to both parental strains (Figure S3), wholesale replacement of recipient SMBGCs by the homologous SMBGCs of the donor was commonly observed. Twenty out of 24 recombinants showed the replacement of one to 14 SMBGCs. These replacements resulted from recombination events occurring in the regions flanking the SMBGCs. Since homologous SMBGCs can differ in gene content (Figure S4), such substitutions potentially result in a significant change in biosynthesis capacity. In a third of the recombinants, a recombination breakpoint occurred within the SMBGC itself resulting in the formation of chimeric SMBGCs (Figure 4). In five out of nine chimeric SMBGCs, the recombination breakpoint(s) occurred in one of the SMBGC biosynthetic genes. This could reflect the recombinogenic nature of SMBGCs *via* sequence repeats associated with redundant protein motifs within multidomain biosynthetic genes^16^. Figure 4 illustrates such cases with recombination in A1 in the NRPS-like/T1PKS (Figure 4a), in recombinant A7 in the aminopolycarboxylic acid biosynthetic gene (Figure 4b) and in recombinant B2 in the siderophore synthase of NI-siderophore (Figure 4c). In the last case (B2), the siderophore SMBGC constitutes the border of a large acquired region, and as depicted in Figure 4c, several recombination breakpoints were identified very close near to each other forming a microheterogeneity hotspot within the siderophore synthase gene.

**Figure 4:**
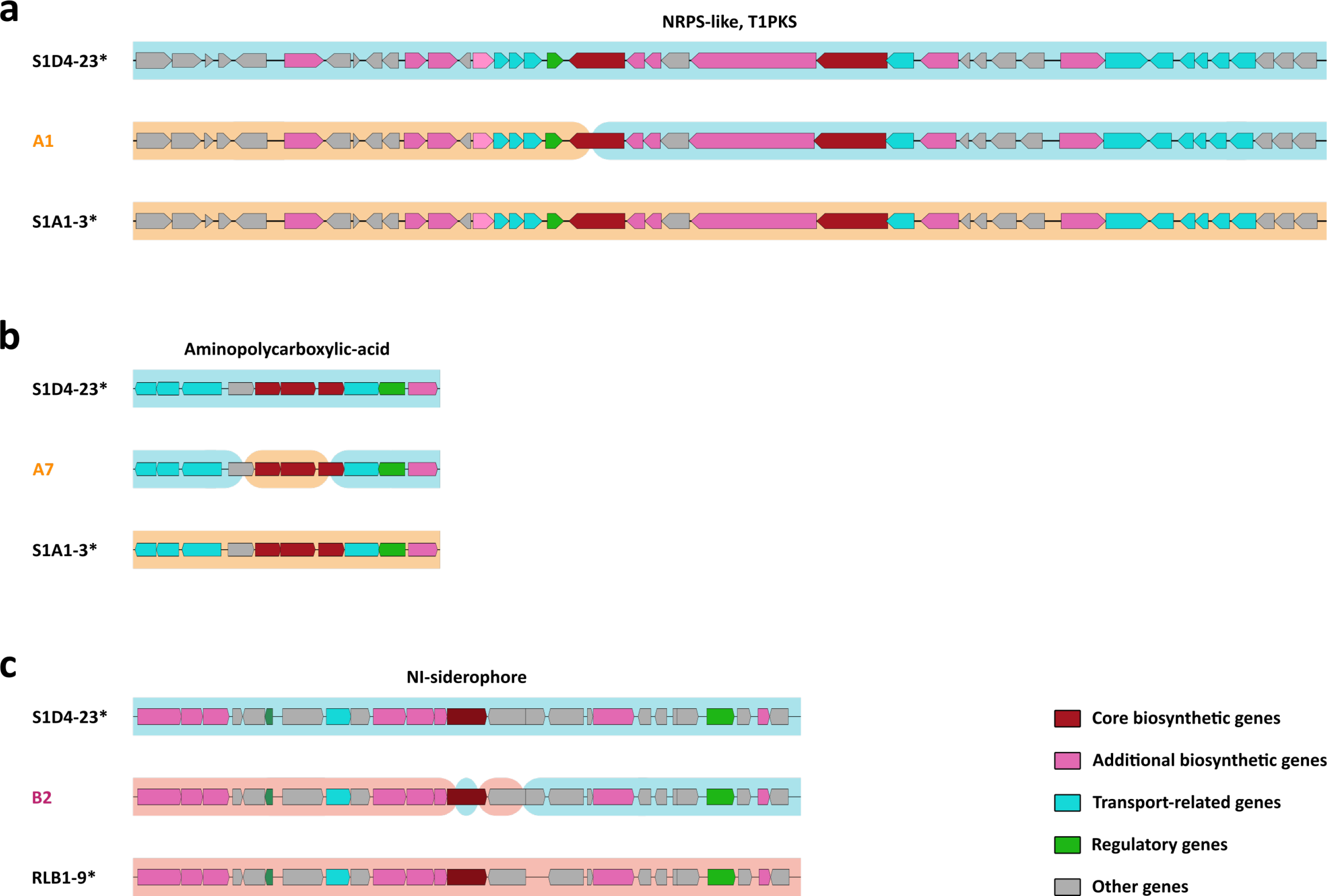
Representation of three different chimeric SMBGCs obtained in the recombinant progeny. Homologous parental SMBGCs predicted by antiSMASH are represented according to this software nomenclature. The donor strain SMBGC is represented at the top of each scheme and the recipient strain SMBGC at the bottom, both SMBGCs are highlighted with a strain specific color according to Figure 1. Chimeric SMBGCs identified in recombinants (A1, A7 and B2) are represented in-between parental SMBGCs. The DNA parental origin in a chimeric SMBGC is identified according to its background color that correspond to those specifically highlighting parental SMBGCs.

**Table 1:** SMBGC reassortment in recombinants. ^#^Replacement of a complete SMBGC in the recipient by its homologue from the donor leads to the formation of a substitution. When the recombination event occurs within the SMBGC, a chimeric version of the cluster is produced.

| Recombinant | A1 | A2 | A3 | A4 | A5 | A6 | A7 | A8 | A9 | A10 | A11 | A12 | B2 | B3 | B4 | B5 | B7 | B8 | B9 | B10 | B11 | B12 | B1 | B6 |
| --- | --- | --- | --- | --- | --- | --- | --- | --- | --- | --- | --- | --- | --- | --- | --- | --- | --- | --- | --- | --- | --- | --- | --- | --- |
| Gain | - | 2 | 2 | 2 | 2 | 2 | 2 | 2 | - | - | - | - | - | - | 2 | - | - | - | - | - | - | - | - | - |
| Loss | 2 | - | 1 | - | - | 2 | - | 4 | - | - | - | - | - | - | 6 | - | - | - | - | - | - | - | - | - |
| Substitution<sup>#</sup> | - | 2 | 6 | 5 | - | 14 | 4 | 5 | - | 2 | - | 6 | 1 | 4 | 2 | 3 | 1 | 3 | 3 | 4 | 3 | 3 | 4 | 2 |
| Chimerization<sup>#</sup> | 1 | - | 1 | - | - | 1 | 2 | 1 | - | - | - | - | 1 | - | - | - | 1 | - | - | - | - | - | 1 | - |

Acquisition of donor-specific SMBGCs by recipients was observed mainly in terminal regions. Two SMBGCs (*terpene* and *Ripp-like*) were acquired in the same DNA fragment in eight recombinants (A2- A8 and B4, Table 1). As the terminal regions are duplicated, all the SMBGCs present in these regions are *de facto* in two copies. Loss of SMBGCs (from one to six) was observed in five recombinants (A1, A3, A6, A8, B4, Table 1). The simplest event corresponded to the replacement of a region harboring a SMBGC in the recipient by its donor homolog devoid of the SMBGC (A3 and A6). In three recombinants (A1, A8 and B4), the partial loss of a SMBGC was not directly linked to recombination with incoming DNA, but to rearrangement of the host genome. In A8, a deletion of about 416 kb resulted in the partial loss of two SMBGCs, but also in the creation of a NRPS and T3PKS hybrid SMBGC at the junction point (Figure S5).

To test whether conjugative transfer can mobilize specialized metabolite biosynthesis pathways, a selection marker (*aac(3)IV* for apramycine resistance) was inserted close to specific SMBGCs in two strains. One marker was inserted close to a SMBGC encoding a lanthipeptide in RLB3-17* (at 80.5 kb), and another close to a SMBGC encoding an NRPS-nucleoside in S1D4-23** (at 27.5 kb). In couple C (RLB3-17* x RLB1-8*, ANI 99.0%), eight recombinants were whole-genome sequenced. Six of them were found to have acquired the lanthipeptide SMBGC. The size of the fragment carrying both the SMBGC and the selection marker varied between 341.9 kb and 609.5 kb. In couple D (S1D4-23** x S1D4-14*, ANI 98.7%), the three recombinants analyzed acquired NRPS-containing fragments ranging in size from 221 kb to 509 kb. The size of recombined chromosomal DNA fragments, several hundred kilobases around the selection marker, is in the same range as that of the SMBGCs, enabling their whole-sale transfer.

## Discussion

Here, we show that AICE transfer in *Streptomyces* is accompanied by the uptake and homologous recombination of vast tracts of unlinked chromosomal DNA. A single conjugation event can result in the replacement of 1.5% to 37.8% of the recipient genome, distributed in fragments across the chromosome: a phenomenon for which we propose the term *Streptomyces* Chromosomal Transfer (SCT). The size and distribution of the fragments indicates that one or more large DNA fragments are transferred followed by recombination with the host chromosome. Microheterogeneity at the edge of large transferred DNA stretches probably result from gene conversion at the recombination site, which is responsible for alternating parental sequences in recombinants^25^. Microheterogeneity hotspots have also been observed in *Mycobacterium*^6^ and *Mycoplasma*^26^, two genera in which largescale HGT events and chromosomal mosaicism have been previously demonstrated^6,26^.

SCT can be hypothesized to occur *via* three distinct TraB-dependent mechanisms (Figure 5). In a first scenario originally put forward by Sir D.A. Hopwood, AICEs transfer without being excised from the donor chromosome^27,28^ (Figure 5a). This hypothesis was proposed based on analysis of chromosome marker mobilization by the integrated form of the conjugative plasmid SCP1 in *Streptomyces coelicolor* A3(2)^8^. Mobilization of the element is here assumed to be based on the recognition of a *cis*-acting pattern, the *clt* site for *c*is-acting *l*ocus of *t*ransfer^29,30^, by the TraB complex (hexamer) forming a conjugation pore from the donor to the recipient cells. The mobilization of the conjugative element from its chromosomal locus, without excision, would here mediate the transfer of chromosomal DNA to one or both sides of the AICE in a mono- or bi-directional pattern, in a similar fashion to the polarized mobilization induced by the F factor integrated into the *E. coli* chromosome (Hfr strain). Such a mechanism of SCT would explain the observation of recombinants formed without concomitant acquisition of autonomous AICE. In a second scenario initially suggested by Pettis and Cohen^31^ and further supported by the work of G. Muth’s group^29^, double-stranded chromosomal DNA is transferred through the TraB pore into the recipient mycelium alongside an independently moving free AICE (Figure 5b). Transfer here is hypothesized to be mediated by *clt*-like sequences distributed through the chromosome (*clc* for *clt*-like chromosomal sequences^29^).

**Figure 5:**
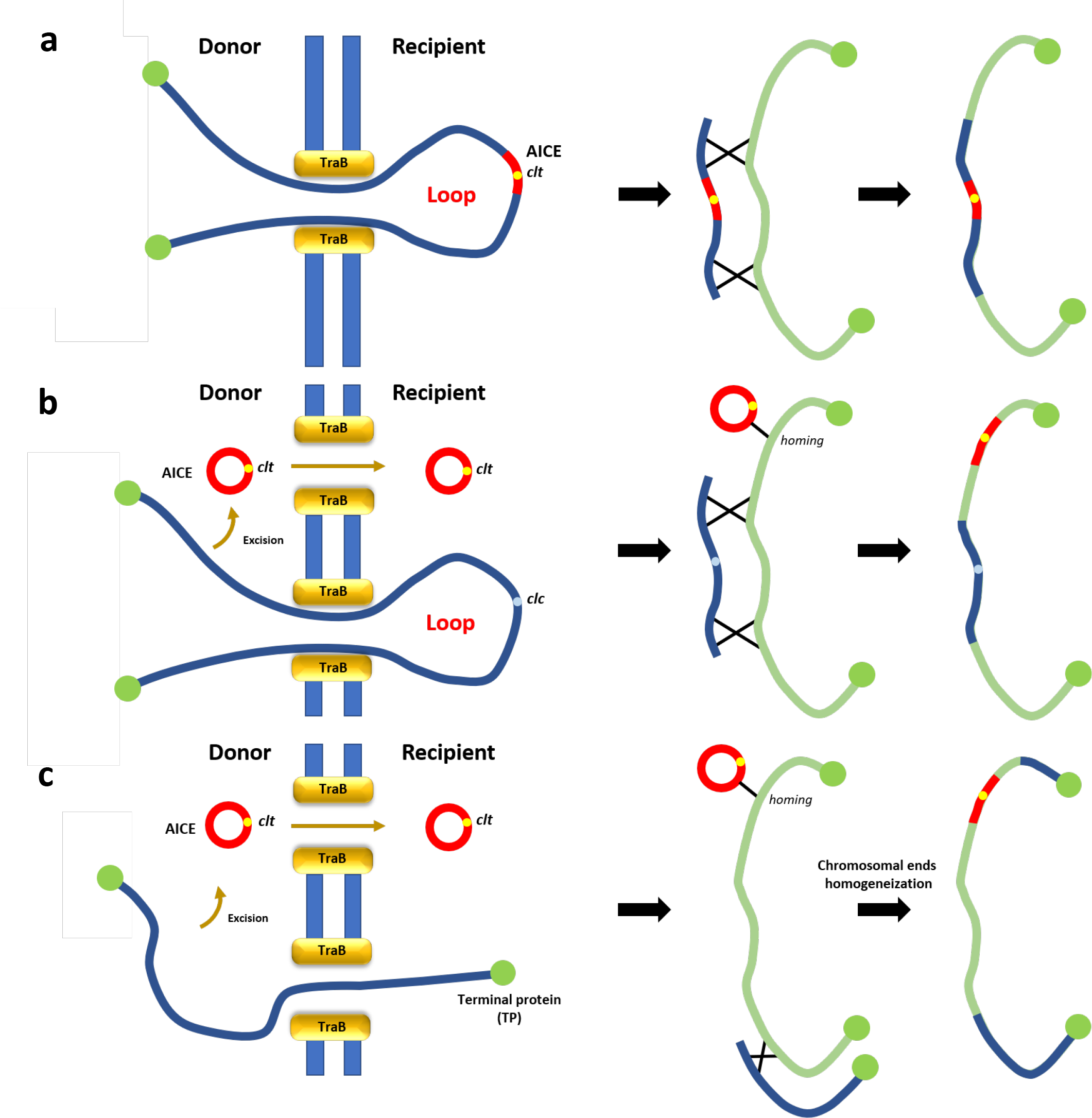
**Scenarios for TraB-dependent DNA mobilization. a**, *cis*-mobilization: mobilization of the conjugative element from its chromosomal locus thanks to its *clt* site for *c*is-acting *l*ocus of *t*ransfer, by the TraB complex (hexamer) forming a conjugation pore from the donor to the recipient cells. A large DNA loop is extruded in the recipient cell. After transfer, replacement of a large region is achieved by homologous recombination. **b**, *trans*-mobilization: chromosomal DNA is transferred through the TraB pore into the recipient mycelium independently from an excised AICE. Transfer of chromosomal DNA is hypothesized to be mediated by *clc* (for *clt*-like chromosomal sequences) distributed through the chromosome. After transfer, self-integration of the AICE occurs at its integration site while chromosomal DNA is integrated by homologous recombination. **c**, ‘End-first’ hypothesis: each chromosomal end could serve as transfer origin and mobilize chromosomal information by the ends. An AICE could self-transfer autonomously. After transfer, the chromosomal end would replace one end and produce a chromosome with two distinct ends; such hybrid structure could be homogenized by terminal recombination. If both ends are transferred and recombined, the concomitant acquisition of the two chromosomal extremities can be obtained. For **b** and **c** mobilization of chromosomal and AICE DNA could occur through a unique TraB pore in the form of double-stranded DNA.

Recombination was frequently accompanied by acquisition of the terminal parts of the chromosome of the donor (seven and one out of twelve in crossing A and B, respectively). The concomitant acquisition of the two chromosomal extremities can be explained by the simultaneous acquisition of chromosome ends. It can also result from the acquisition of a copy of a chromosome arm, the transient production of a hybrid chromosome with divergent TIRs, followed by homogenization of the ends *via* homologous recombination. This homogenization phenomenon is possible thanks to the strong DNA identity between the parental TIRs in their internal parts (between 147 kb and 188 kb from the start of the TIRs of the parental strains are homologous for couple A and B respectively^24^); the more distal region till the chromosome extremity is specific of each strain do not support homologous recombination. A close look at the origin of the SNPs of the terminal regions of the recombinant revealed that frequent exchanges occurred between the homologous parts of the parental TIRs supporting a homogenization mechanism powered by homologous recombination. A third scenario is supported by our recombinant analysis, the so-called ‘end-first’ model proposed by C.W. Chen^32,33^ where chromosomal ends could serve as transfer origins (Figure 5c). In these experiments mobilization of the chromosomal ends was associated with the presence of conjugative plasmids. In our experiments, parental strains of couple A were devoid of conjugative plasmids showing that, in this scenario, AICEs would also be able to promote this end-first mobilization. As all three types of recombination were observed within the population, or even within a single recombinant, the three mechanisms are not mutually exclusive and could operate simultaneously. *Streptomyces* are well-known for their production of antibiotics and other specialized metabolites^34^. The biotechnological exploitation of this biosynthetic arsenal is a key objective in the arms race against the emergence of antibiotic resistance in pathogenic bacteria^35–37^. Cloning and transfer of an entire biosynthetic pathway (often approaching 100 kb in size) is often a hindrance to heterologous expression approaches to identify the metabolite encoded by the SMBGC^38–41^. Our mating assays demonstrated that conjugative transfer resulted in the transfer of complete SMBGCs as well as the shuffling of parental SMBGCs within recombinants. Gain, loss, duplication and (wholesale) SMBGC replacement was observed, especially in terminal regions which are known to be enriched in SMBGCs^42^. To our knowledge, this is the first experimental demonstration of chromosomal SMBGC transfer and recombination in *Streptomyces*.

Recombination within SMBGCs (*i.e.* formation of chimeric and hybrid SMBGCs) was also frequently observed, which could be aided by their repetitive structure^16,43^. PKS and NRPS synthases are large multidomain enzymes encoded by long highly repetitive DNA sequences that can also exhibit short repeated DNA motifs^43^, favouring recombination and gene conversion^16^. The potential of SMBGCs to evolve rapidly might aid adaptation to the competitive and changing environment of the soil. The rapid generation of SMBGC diversification in simple mating type assays also offers a new avenue into natural product discovery by providing new combinations of biosynthesis genes, reassociation of functional domains within SMBGCs and identification of new molecules of interest.

## Material and Methods

### Bacterial strains

This study employed six closely related *Streptomyces* strains that were isolated at a microscale from forest soil in a previous study^19^. To select for recombinants, each of the parental strains was labeled with a specific resistant gene (either *neo* for kanamycin or *aac(3)IV* for apramycin resistance) inserted at different chromosomal positions corresponding to intergenic regions present between two convergent genes (strains *^/^**, Table S1). Briefly, resistance genes were amplified by PCR from the SuperCos1 (Agilent Technologies) and pIJ6902^44^ plasmids respectively (Agilent, USA) (Table S3). Upstream and downstream regions of the targeted chromosomal site were amplified and then assembled by overlap extension-PCR^45^ and cloned (site *Hin*dIII) into the derived pWED2 suicide vector (pWED2*^11^). The *neo* or *aac(3)IV* genes were then inserted and cloned into a *Xba*I site designed in for that purpose between the cloned upstream and downstream regions. These constructs were introduced into *Streptomyces* by intergeneric conjugation with *E. coli* ET12567/pUZ8002^46^ and allelic replacement was selected on antibiotic resistance (*neo* or *aac(3)IV*) and susceptibility to vector-borne resistance. Alternatively, strains labeled by simple crossover integration of the construct in the chromosome (via recombination with the upstream or downstream region only) were also used (Table S1). For strain S1D4-23** (S1D4-23 NRPSaac(3)IV), a region close to the SMBGC NRPS nucleoside was amplified by PCR with primers NRPSBamHI_F and NRPSBamHI_R (Table S3), cloned into the suicide vector pIJ8668^47^. Integration into the chromosome was selected on the basis of the plasmid-borne resistance (apramycin). The labeled strains are listed in Table S1. In order to isolate recombinants, conjugations were performed with approximately 10^6^ spores of each labeled parental strain cocultured on Hickey-Tresner (HT) agar medium^48^ for 7 days at 30°C. The bacterial lawn was replicated on Mannitol Soy Flour medium (SFM) agar^46^ supplemented with 50 µg/ml kanamycin and 50 µg/ml apramycin, and grown for 5 days at 30°C. The double-resistant colonies were then subcloned twice to ensure genetic homogeneity. The ratio of the number of double-resistant colonies to the total number of spores (both parental strains can be donor) was used to calculate recombination frequency.

### Genome sequencing

Genomic DNA extraction was performed according to Kieser et al.^46^. The genome of recombinants was sequenced by MiSeq sequencing (Illumina, CA, USA) (mean coverage 100x) at the Plateforme de Microbiologie Mutualisée (P2M) of the Pasteur Institute (Paris). The genome data are available on the NCBI database for the parental strains (Table S1) and raw sequencing data for the recombinant strains were deposited at SRA (NCBI, Bioproject PRJNA912173).

### Analysis of recombinant genomes

For each recombinant genome, sequencing reads were aligned with both parental strains to identify recipient and donor parent genome. The alignments were performed using the Geneious prime platform reference algorithm (Invitrogen Corp., version 2022.0.1) with the following parameters: 2 iterations, minimum quality mapping 30, minimum coverage 20x (if this threshold was not reached Ns were added). The consensus of the recombinant sequence was aligned with the chromosomal sequences of the parents using the progressive MAUVE algorithm^49^. The Single Nucleotide Polymorphism (SNP) call table was exported with MAUVE tool ‘export SNPs’. Custom scripts (Python V.3.9) were used to call SNPs of both parental strains along the alignment (Github). The presence of two or more successive SNPs was used to assign a genomic region to one of the two parents (to buffer against possible effects of - single - point mutations). The parental origin of the TIRs of each recombinant could be unambiguously assigned thanks to the 116-220 kb to 246-261 kb which are specific to the parental chromosomes at the very ends of the TIRs in couple A and B, respectively. The output of analyses was manually checked to eliminate artifacts related to repeated sequences such as insertion sequences. Specialized Metabolite Biosynthetic Gene Clusters (SMBGCs) were predicted using antiSMASH^50^. Whole chromosome structure of parental and recombinant strains was visualized in a circular layout produced by Circos^51^.

### Data availability

Raw sequencing data for the recombinant strains were deposited at SRA (NCBI, Bioproject PRJNA912173).

### Code availability

The codes to reproduce the main results have been deposited on GitHub: https://github.com/leblond14u/Large-scale-chromosomal-DNA-transfer

## Acknowledgements

We would like to thank Jean-Luc Pernodet (I2BC, CNRS) for his valuable advice on improving the manuscript.

