## Supplementary figures and tables for "Conjugation mediates large-scale chromosomal transfer in *Streptomyces* driving diversification of antibiotic biosynthetic gene clusters": Supp_data_Choufa_27052024.pptx

### Slide 1
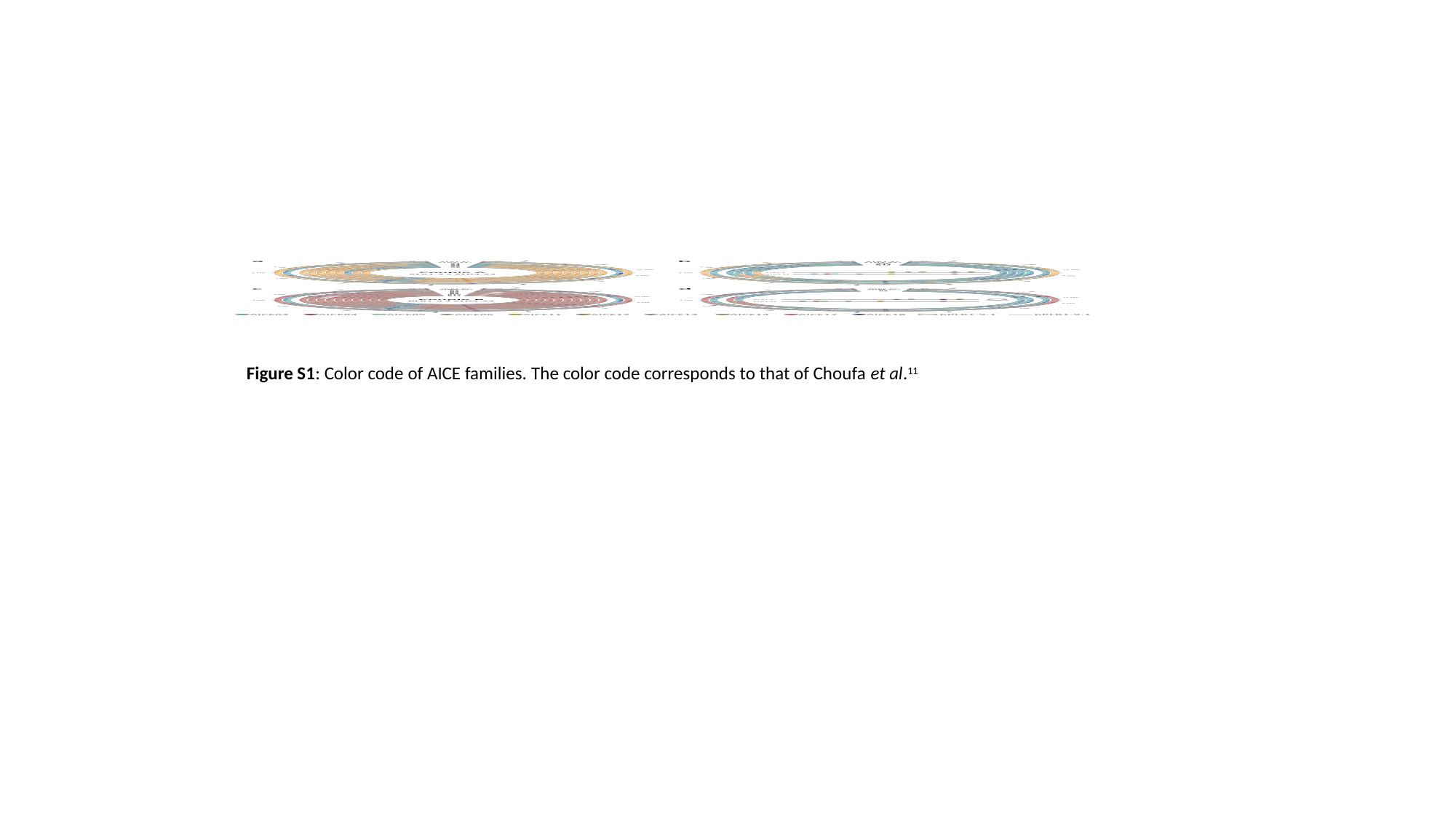

Figure S1: Color code of AICE families. The color code corresponds to that of Choufa et al.^11^

### Slide 2
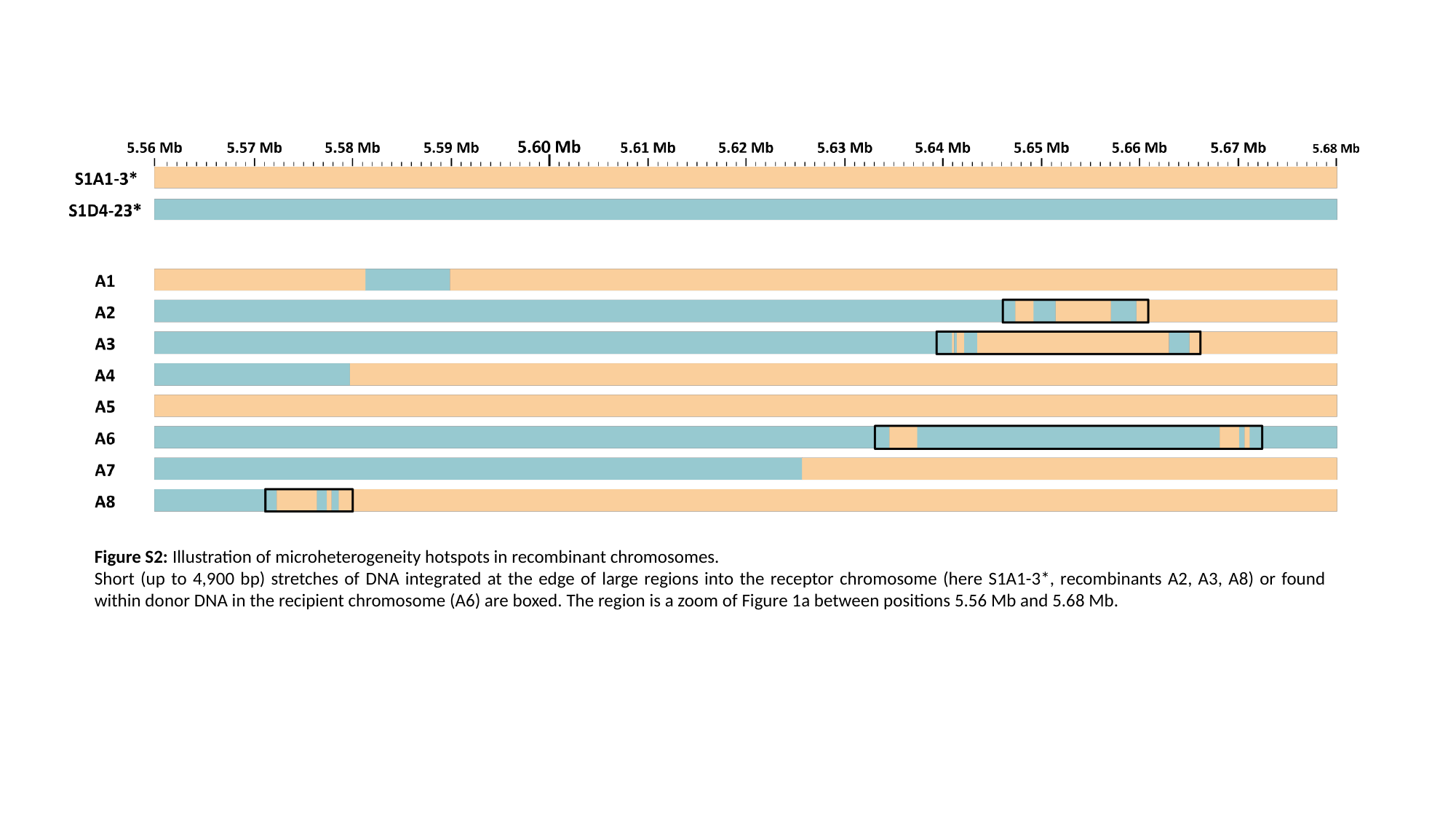

Figure S2: Illustration of microheterogeneity hotspots in recombinant chromosomes.
Short (up to 4,900 bp) stretches of DNA integrated at the edge of large regions into the receptor chromosome (here S1A1-3*, recombinants A2, A3, A8) or found within donor DNA in the recipient chromosome (A6) are boxed. The region is a zoom of Figure 1a between positions 5.56 Mb and 5.68 Mb.

### Slide 3
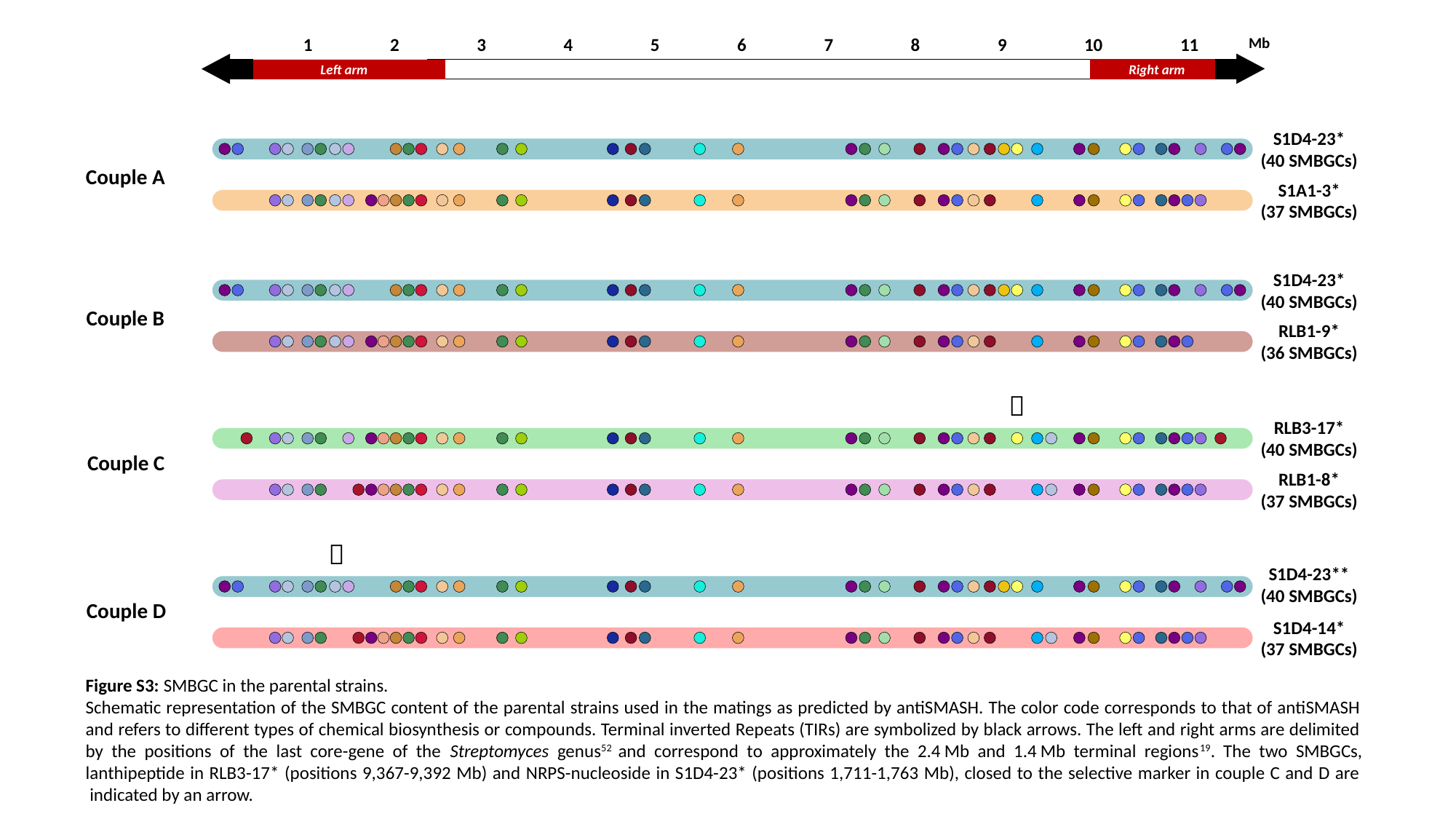

1
2
3
4
5
6
7
8
9
10
11
Mb
Right arm
Left arm
S1D4-23*
(40 SMBGCs)
Couple A
S1A1-3*
(37 SMBGCs)
S1D4-23*
(40 SMBGCs)
Couple B
RLB1-9*
(36 SMBGCs)
*
RLB3-17*
(40 SMBGCs)
Couple C
RLB1-8*
(37 SMBGCs)
*
S1D4-23**
(40 SMBGCs)
Couple D
S1D4-14*
(37 SMBGCs)
🠛
🠛
Figure S3: SMBGC in the parental strains.
Schematic representation of the SMBGC content of the parental strains used in the matings as predicted by antiSMASH. The color code corresponds to that of antiSMASH and refers to different types of chemical biosynthesis or compounds. Terminal inverted Repeats (TIRs) are symbolized by black arrows. The left and right arms are delimited by the positions of the last core-gene of the Streptomyces genus^52^ and correspond to approximately the 2.4 Mb and 1.4 Mb terminal regions^19^. The two SMBGCs, lanthipeptide in RLB3-17* (positions 9,367-9,392 Mb) and NRPS-nucleoside in S1D4-23* (positions 1,711-1,763 Mb), closed to the selective marker in couple C and D are  indicated by an arrow.

### Slide 4
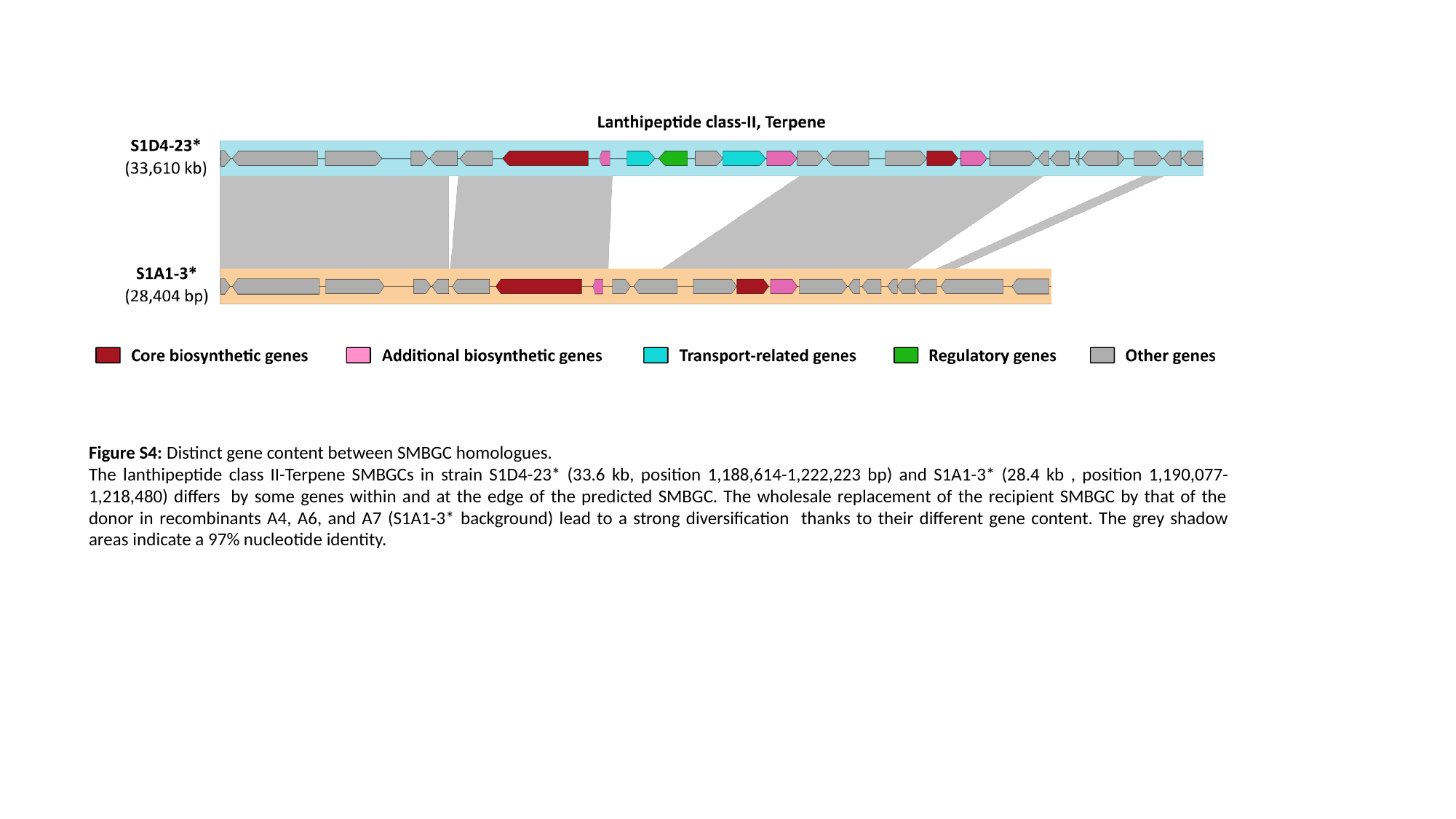

Figure S4: Distinct gene content between SMBGC homologues.
The lanthipeptide class II-Terpene SMBGCs in strain S1D4-23* (33.6 kb, position 1,188,614-1,222,223 bp) and S1A1-3* (28.4 kb , position 1,190,077-1,218,480) differs  by some genes within and at the edge of the predicted SMBGC. The wholesale replacement of the recipient SMBGC by that of the donor in recombinants A4, A6, and A7 (S1A1-3* background) lead to a strong diversification  thanks to their different gene content. The grey shadow areas indicate a 97% nucleotide identity.

### Slide 5
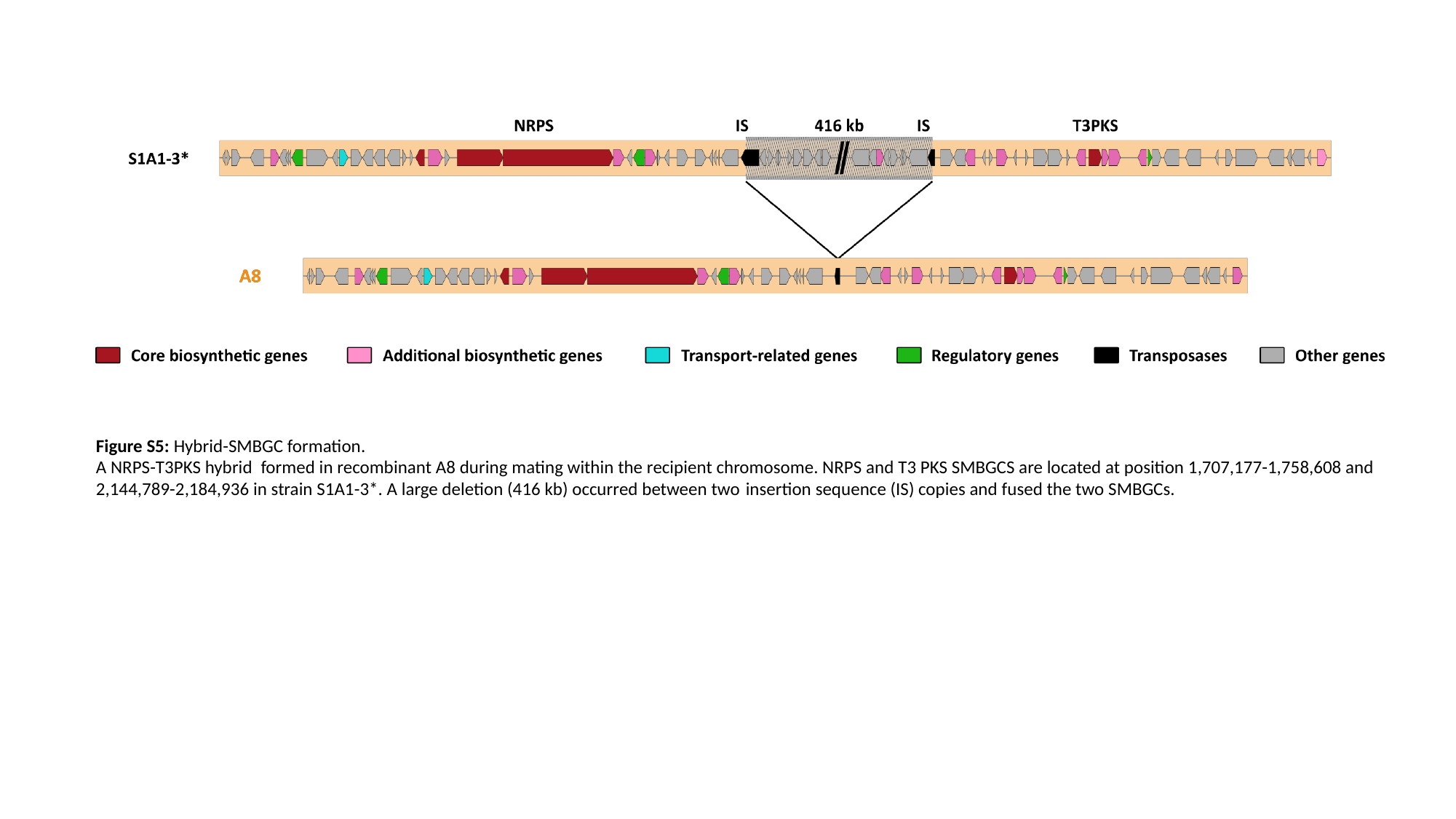

Figure S5: Hybrid-SMBGC formation.
A NRPS-T3PKS hybrid  formed in recombinant A8 during mating within the recipient chromosome. NRPS and T3 PKS SMBGCS are located at position 1,707,177-1,758,608 and 2,144,789-2,184,936 in strain S1A1-3*. A large deletion (416 kb) occurred between two insertion sequence (IS) copies and fused the two SMBGCs.

### Slide 6
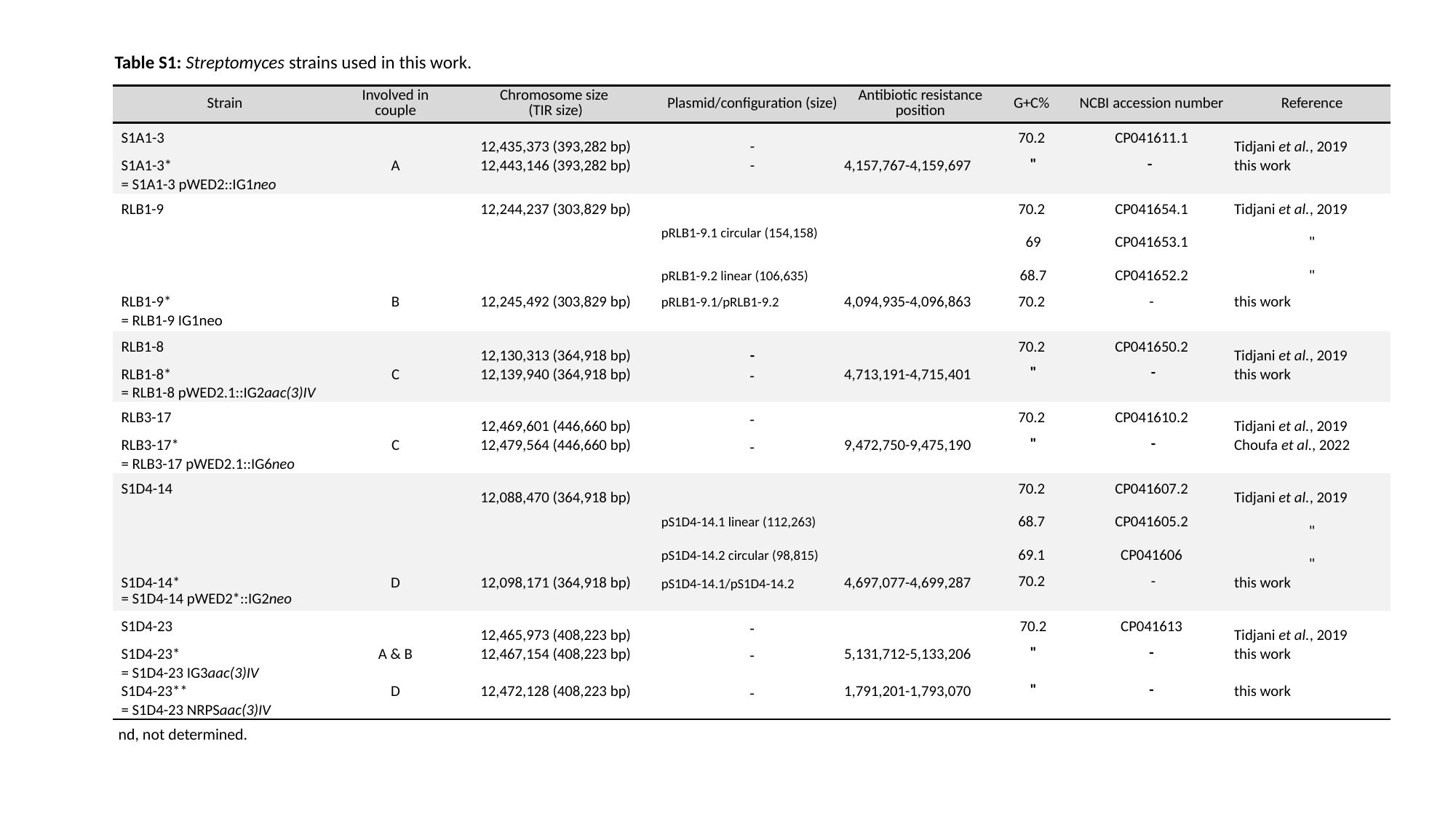

Table S1: Streptomyces strains used in this work.
| Strain | Involved in couple | Chromosome size (TIR size) | Plasmid/configuration (size) | Antibiotic resistance position | G+C% | NCBI accession number | Reference |
| --- | --- | --- | --- | --- | --- | --- | --- |
| S1A1-3 | | 12,435,373 (393,282 bp) | - | | 70.2 | CP041611.1 | Tidjani et al., 2019 |
| S1A1-3\* | A | 12,443,146 (393,282 bp) | - | 4,157,767-4,159,697 | " | - | this work |
| = S1A1-3 pWED2::IG1neo | | | | | | | |
| RLB1-9 | | 12,244,237 (303,829 bp) | | | 70.2 | CP041654.1 | Tidjani et al., 2019 |
| | | | pRLB1-9.1 circular (154,158) | | 69 | CP041653.1 | " |
| | | | pRLB1-9.2 linear (106,635) | | 68.7 | CP041652.2 | " |
| RLB1-9\* | B | 12,245,492 (303,829 bp) | pRLB1-9.1/pRLB1-9.2 | 4,094,935-4,096,863 | 70.2 | - | this work |
| = RLB1-9 IG1neo | | | | | | | |
| RLB1-8 | | 12,130,313 (364,918 bp) | - | | 70.2 | CP041650.2 | Tidjani et al., 2019 |
| RLB1-8\* | C | 12,139,940 (364,918 bp) | - | 4,713,191-4,715,401 | " | - | this work |
| = RLB1-8 pWED2.1::IG2aac(3)IV | | | | | | | |
| RLB3-17 | | 12,469,601 (446,660 bp) | - | | 70.2 | CP041610.2 | Tidjani et al., 2019 |
| RLB3-17\* | C | 12,479,564 (446,660 bp) | - | 9,472,750-9,475,190 | " | - | Choufa et al., 2022 |
| = RLB3-17 pWED2.1::IG6neo | | | | | | | |
| S1D4-14 | | 12,088,470 (364,918 bp) | | | 70.2 | CP041607.2 | Tidjani et al., 2019 |
| | | | pS1D4-14.1 linear (112,263) | | 68.7 | CP041605.2 | " |
| | | | pS1D4-14.2 circular (98,815) | | 69.1 | CP041606 | " |
| S1D4-14\* | D | 12,098,171 (364,918 bp) | pS1D4-14.1/pS1D4-14.2 | 4,697,077-4,699,287 | 70.2 | - | this work |
| = S1D4-14 pWED2\*::IG2neo | | | | | | | |
| S1D4-23 | | 12,465,973 (408,223 bp) | - | | 70.2 | CP041613 | Tidjani et al., 2019 |
| S1D4-23\* | A & B | 12,467,154 (408,223 bp) | - | 5,131,712-5,133,206 | " | - | this work |
| = S1D4-23 IG3aac(3)IV | | | | 1,791,201-1,793,070 | | | |
| S1D4-23\*\* | D | 12,472,128 (408,223 bp) | - | | " | - | this work |
| = S1D4-23 NRPSaac(3)IV | | | | | | | |
nd, not determined.

### Slide 7
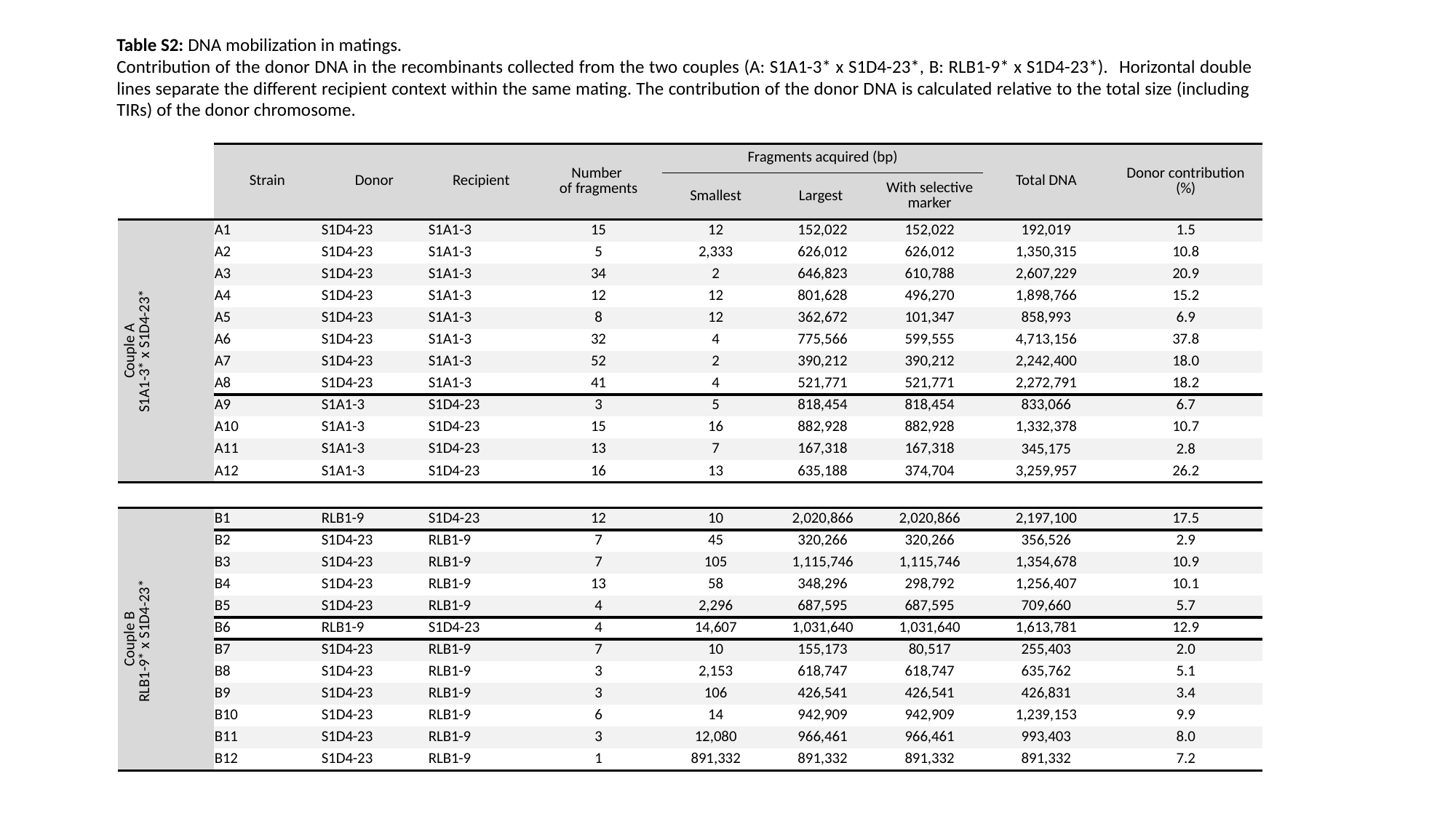

Table S2: DNA mobilization in matings.
Contribution of the donor DNA in the recombinants collected from the two couples (A: S1A1-3* x S1D4-23*, B: RLB1-9* x S1D4-23*).  Horizontal double lines separate the different recipient context within the same mating. The contribution of the donor DNA is calculated relative to the total size (including TIRs) of the donor chromosome.
| | Strain | Donor | Recipient | Number of fragments | Fragments acquired (bp) | | | Total DNA | Donor contribution (%) |
| --- | --- | --- | --- | --- | --- | --- | --- | --- | --- |
| | | | | | Smallest | Largest | With selective marker | | |
| Couple A S1A1-3\* x S1D4-23\* | A1 | S1D4-23 | S1A1-3 | 15 | 12 | 152,022 | 152,022 | 192,019 | 1.5 |
| | A2 | S1D4-23 | S1A1-3 | 5 | 2,333 | 626,012 | 626,012 | 1,350,315 | 10.8 |
| | A3 | S1D4-23 | S1A1-3 | 34 | 2 | 646,823 | 610,788 | 2,607,229 | 20.9 |
| | A4 | S1D4-23 | S1A1-3 | 12 | 12 | 801,628 | 496,270 | 1,898,766 | 15.2 |
| | A5 | S1D4-23 | S1A1-3 | 8 | 12 | 362,672 | 101,347 | 858,993 | 6.9 |
| | A6 | S1D4-23 | S1A1-3 | 32 | 4 | 775,566 | 599,555 | 4,713,156 | 37.8 |
| | A7 | S1D4-23 | S1A1-3 | 52 | 2 | 390,212 | 390,212 | 2,242,400 | 18.0 |
| | A8 | S1D4-23 | S1A1-3 | 41 | 4 | 521,771 | 521,771 | 2,272,791 | 18.2 |
| | A9 | S1A1-3 | S1D4-23 | 3 | 5 | 818,454 | 818,454 | 833,066 | 6.7 |
| | A10 | S1A1-3 | S1D4-23 | 15 | 16 | 882,928 | 882,928 | 1,332,378 | 10.7 |
| | A11 | S1A1-3 | S1D4-23 | 13 | 7 | 167,318 | 167,318 | 345,175 | 2.8 |
| | A12 | S1A1-3 | S1D4-23 | 16 | 13 | 635,188 | 374,704 | 3,259,957 | 26.2 |
| Couple B RLB1-9\* x S1D4-23\* | B1 | RLB1-9 | S1D4-23 | 12 | 10 | 2,020,866 | 2,020,866 | 2,197,100 | 17.5 |
| | B2 | S1D4-23 | RLB1-9 | 7 | 45 | 320,266 | 320,266 | 356,526 | 2.9 |
| | B3 | S1D4-23 | RLB1-9 | 7 | 105 | 1,115,746 | 1,115,746 | 1,354,678 | 10.9 |
| | B4 | S1D4-23 | RLB1-9 | 13 | 58 | 348,296 | 298,792 | 1,256,407 | 10.1 |
| | B5 | S1D4-23 | RLB1-9 | 4 | 2,296 | 687,595 | 687,595 | 709,660 | 5.7 |
| | B6 | RLB1-9 | S1D4-23 | 4 | 14,607 | 1,031,640 | 1,031,640 | 1,613,781 | 12.9 |
| | B7 | S1D4-23 | RLB1-9 | 7 | 10 | 155,173 | 80,517 | 255,403 | 2.0 |
| | B8 | S1D4-23 | RLB1-9 | 3 | 2,153 | 618,747 | 618,747 | 635,762 | 5.1 |
| | B9 | S1D4-23 | RLB1-9 | 3 | 106 | 426,541 | 426,541 | 426,831 | 3.4 |
| | B10 | S1D4-23 | RLB1-9 | 6 | 14 | 942,909 | 942,909 | 1,239,153 | 9.9 |
| | B11 | S1D4-23 | RLB1-9 | 3 | 12,080 | 966,461 | 966,461 | 993,403 | 8.0 |
| | B12 | S1D4-23 | RLB1-9 | 1 | 891,332 | 891,332 | 891,332 | 891,332 | 7.2 |

### Slide 8
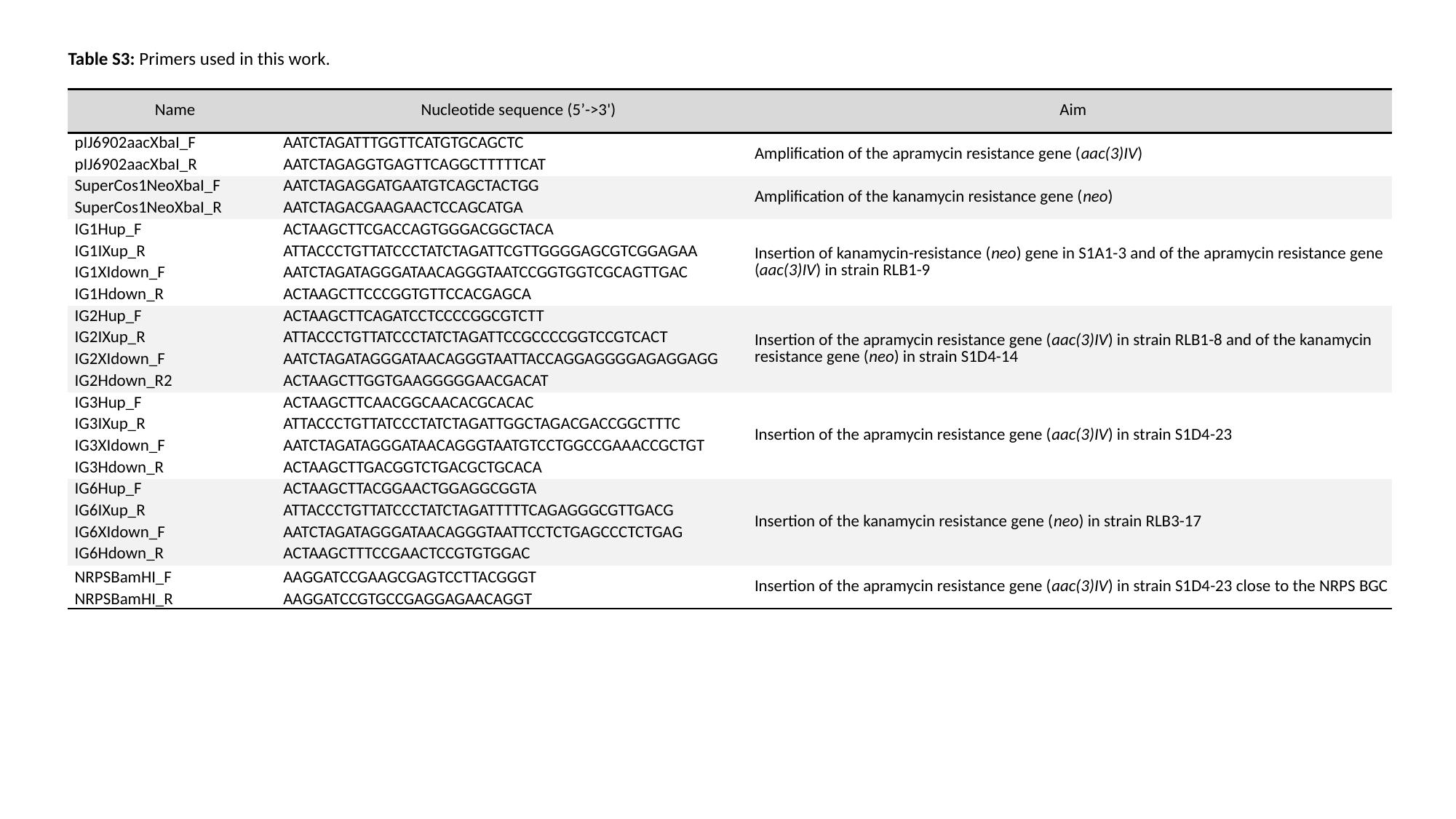

Table S3: Primers used in this work.
| Name | Nucleotide sequence (5’->3') | Aim |
| --- | --- | --- |
| pIJ6902aacXbaI\_F | AATCTAGATTTGGTTCATGTGCAGCTC | Amplification of the apramycin resistance gene (aac(3)IV) |
| pIJ6902aacXbaI\_R | AATCTAGAGGTGAGTTCAGGCTTTTTCAT | |
| SuperCos1NeoXbaI\_F | AATCTAGAGGATGAATGTCAGCTACTGG | Amplification of the kanamycin resistance gene (neo) |
| SuperCos1NeoXbaI\_R | AATCTAGACGAAGAACTCCAGCATGA | |
| IG1Hup\_F | ACTAAGCTTCGACCAGTGGGACGGCTACA | Insertion of kanamycin-resistance (neo) gene in S1A1-3 and of the apramycin resistance gene (aac(3)IV) in strain RLB1-9 |
| IG1IXup\_R | ATTACCCTGTTATCCCTATCTAGATTCGTTGGGGAGCGTCGGAGAA | |
| IG1XIdown\_F | AATCTAGATAGGGATAACAGGGTAATCCGGTGGTCGCAGTTGAC | |
| IG1Hdown\_R | ACTAAGCTTCCCGGTGTTCCACGAGCA | |
| IG2Hup\_F | ACTAAGCTTCAGATCCTCCCCGGCGTCTT | Insertion of the apramycin resistance gene (aac(3)IV) in strain RLB1-8 and of the kanamycin resistance gene (neo) in strain S1D4-14 |
| IG2IXup\_R | ATTACCCTGTTATCCCTATCTAGATTCCGCCCCGGTCCGTCACT | |
| IG2XIdown\_F | AATCTAGATAGGGATAACAGGGTAATTACCAGGAGGGGAGAGGAGG | |
| IG2Hdown\_R2 | ACTAAGCTTGGTGAAGGGGGAACGACAT | |
| IG3Hup\_F | ACTAAGCTTCAACGGCAACACGCACAC | Insertion of the apramycin resistance gene (aac(3)IV) in strain S1D4-23 |
| IG3IXup\_R | ATTACCCTGTTATCCCTATCTAGATTGGCTAGACGACCGGCTTTC | |
| IG3XIdown\_F | AATCTAGATAGGGATAACAGGGTAATGTCCTGGCCGAAACCGCTGT | |
| IG3Hdown\_R | ACTAAGCTTGACGGTCTGACGCTGCACA | |
| IG6Hup\_F | ACTAAGCTTACGGAACTGGAGGCGGTA | Insertion of the kanamycin resistance gene (neo) in strain RLB3-17 |
| IG6IXup\_R | ATTACCCTGTTATCCCTATCTAGATTTTTCAGAGGGCGTTGACG | |
| IG6XIdown\_F | AATCTAGATAGGGATAACAGGGTAATTCCTCTGAGCCCTCTGAG | |
| IG6Hdown\_R | ACTAAGCTTTCCGAACTCCGTGTGGAC | |
| NRPSBamHI\_F | AAGGATCCGAAGCGAGTCCTTACGGGT | Insertion of the apramycin resistance gene (aac(3)IV) in strain S1D4-23 close to the NRPS BGC |
| NRPSBamHI\_R | AAGGATCCGTGCCGAGGAGAACAGGT | |
